# EvoSynth: Enabling Multi-Target Drug Discovery through Latent Evolutionary Optimization and Synthesis-Aware Prioritization

**DOI:** 10.1101/2025.11.04.686584

**Authors:** Viet Thanh Duy Nguyen, Phuc Pham, Truong-Son Hy

**Affiliations:** Department of Computer Science, the University of Alabama at Birmingham, Alabama, United States

## Abstract

Complex diseases, such as cancer and neurodegeneration, feature interconnected pathways, making single-target therapies ineffective due to pathway redundancy and compensatory mechanisms. Polypharmacy, which combines multiple drugs to target distinct proteins, addresses this but often leads to drug-drug interactions, cumulative toxicity, and complex pharmacokinetics. To overcome these challenges, we introduce EvoSynth, a modular framework for multi-target drug discovery that combines latent evolution and synthesis-aware prioritization to generate and prioritize candidates with high translational potential. Latent evolution navigates a chemically and functionally informed latent space to identify candidates with strong predicted affinity across multiple targets. Synthesis-aware prioritization evaluates both retrosynthetic feasibility and the trade-off between synthetic cost and therapeutic reward, enabling practical and efficient candidate selection. Applied to dual inhibition of JNK3 and GSK3*β* in Alzheimer’s disease and PI3K and PARP1 in ovarian cancer, EvoSynth consistently outperforms baseline generative models, achieving higher predicted affinities, improved scaffold diversity, and lower synthesis costs. These findings highlight EvoSynth’s ability to integrate target-driven generation with practical synthesizability, establishing a scalable framework for multi-target and polypharmacological drug discovery. Our source code and data to reproduce all experiments is publicly available on GitHub at: https://github.com/HySonLab/EvoSynth.

## 1 Introduction

Historically, drug discovery has emphasized high selectivity toward a single biological target to minimize off-target effects and reduce the risk of adverse reactions. As a result, compounds exhibiting activity against multiple proteins were traditionally viewed as undesirable. However, this single-target paradigm has proven inadequate for treating complex, multifactorial diseases, which often involve dysregulation across interconnected pathways^1^. Alternatively, polypharmacy, the use of multiple drugs to target distinct proteins, has been adopted to address the complexity of such diseases. Although this approach can improve therapeutic coverage, it often leads to drug-drug interactions, increased toxicity, inconsistent pharmacokinetics, and reduced patient adherence due to complex dosing regimens^2^. Multi-target drug discovery has emerged as a promising paradigm for addressing the limitations of both single-target therapies and polypharmacy. By designing compounds capable of selectively interacting with multiple disease-relevant targets, this approach aims to restore disrupted network dynamics and produce additive or synergistic therapeutic effects. It is particularly attractive for complex diseases, where the simultaneous modulation of interconnected pathways is often required to achieve durable clinical responses^3^. This strategy has already shown encouraging results in several well-known multifactorial diseases, including neurodegenerative diseases^4^ (such as Alzheimer’s and Parkinson’s disease) and cancers^5^ (such as breast, lung, and ovarian cancer), further validating its therapeutic potential.

Recent advances in artificial intelligence (AI) have revolutionized early-stage drug discovery, enabling rapid molecular generation, property prediction, and structure-based design. However, most AI-driven frameworks remain tailored to single-target settings^6^, limiting their utility for multifactorial diseases. A growing number of efforts have begun to adapt AI for multi-target applications. One notable example is MolSculptor^7^, which integrates latent diffusion with evolutionary optimization to generate multi-site inhibitors without requiring extensive task-specific training data. Despite promising results, MolSculptor exhibits several limitations that constrain its applicability in real-world drug discovery pipelines. First, its latent space is intentionally trained for structural reconstruction to allow for a broad generalization without requiring extensive task-specific data. Although this design facilitates flexible multi-site inhibitor generation, it lacks explicit encoding of functional properties such as binding affinity, which limits the model’s ability to prioritize candidates with strong multi-target bioactivity. Second, while MolSculptor employs empirical constraints such as synthetic accessibility (SA)^8^ and quantitative estimation of drug-likeness (QED)^9^, these overlook more clinically relevant ADMET properties, including toxicity, metabolic stability, and permeability of the blood-brain barrier (BBB). These omissions reduce the translational relevance of its output.

A critical limitation of many AI-driven drug design frameworks is their inability to account for the synthesizability of the molecules they generate. Most models are optimized for predicted biological activity, yet rarely assess whether proposed compounds can be synthesized in practice or at what cost. As a result, many promising candidates that appear in silico prove to be infeasible in laboratory settings, limiting their translational value. Two main strategies have emerged to incorporate synthesizability into the design process. Bottom-up approaches use fragment-based generation, assembling molecules from predefined building blocks or reaction templates to ensure synthetic feasibility^10,11^. Although this strategy can in principle yield synthesizable outputs, it is constrained by the choice of fragments and assembly rules, which limits chemical diversity. As the number of functional groups and motifs grows, the combinatorial space expands exponentially, quickly becoming intractable. These approaches also require extensive motif-level annotations, which are often impractical and resource-intensive. Top-down approaches, in contrast, apply retrosynthetic planning to fully generated molecules, assessing their feasibility independently of the generative process. This paradigm is more flexible, as it can be applied to the output of any model but is typically treated as a post hoc filtering step rather than being integrated directly into molecular design. Substantial progress has been made in advancing top-down synthesis planning and decision-support tools. Platforms such as ASKCOS^12^ and AiZynthFinder^13^ enable automated prediction of retrosynthetic routes, while frameworks such as SPARROW^14^ provide principled strategies to prioritize compounds based on the trade-off between therapeutic reward and synthetic cost. However, to the best of our knowledge, few generative drug design frameworks integrate these capabilities directly into the design pipeline. Consequently, the responsibility for assessing the synthesizability and prioritizing candidates often remains disconnected from the generative process, undermining the translation of AI-designed molecules into actionable experimental candidates.

In this work, we introduce EvoSynth, a modular framework for multi-target drug discovery that addresses current limitations by integrating latent evolutionary optimization, biological relevance, and synthesis-aware prioritization into a unified design pipeline. EvoSynth performs evolutionary optimization in a score-informed latent space that encodes both structural and functional information, enabling the search to efficiently converge on candidates with strong predicted affinity in multiple targets. To ensure clinical relevance, we incorporate a comprehensive set of pharmacological property filters, including SA, QED and ADMET-related constraints that are tailored to the therapeutic context, such as the permeability of the BBB for Alzheimer’s disease and the toxicity profiles for OC. For synthesis-aware prioritization, we adopted the SPARROW framework, which evaluates retrosynthetic feasibility while optimizing the trade-off between synthetic cost and therapeutic reward. We argue that SPARROW is particularly well suited to EvoSynth’s evolutionary sampling strategy, which naturally produces families of structurally related molecules, increasing the likelihood of shared intermediates and overlapping synthetic routes. This synergy enables optimization of batch synthesis under realistic resource constraints. We demonstrate the utility of EvoSynth in dual-target scenarios in Alzheimer’s disease (JNK3 and GSK3*β*)^15^ and ovarian cancer (PI3K and PARP1)^16^, showing that it consistently generates druglike molecules with robust dual-target activity and tractable synthesis routes.

## 2 Results

We evaluated the effectiveness of EvoSynth in the context of dual-target drug discovery by comparing it with MolSculptor^7^, the most closely related framework that combines latent diffusion with evolutionary optimization for the generation of multi-site inhibitors. As EvoSynth builds on the core architecture and guiding principles of MolSculptor, this comparison allows us to directly assess the benefits of our proposed methodological enhancements. To this end, we consider dual-target tasks of high clinical significance, initializing each with a compound that exhibits experimentally validated activity against at least one of the intended targets. This scaffold-based initialization anchors the search in a chemically plausible space and facilitates the generation of optimized analogues with desired polypharmacological profiles. The following case studies illustrate the application of EvoSynth in two distinct therapeutic settings.

- **Alzheimer’s disease (AD):** A leading cause of dementia and a major global health burden^17^. AD is a multifactorial neurodegenerative disorder that involves amyloid-*β* aggregation, tau hyperphosphorylation, oxidative stress, and neuroinflammation^18^. Potential targets, N-terminal kinase 3 (JNK3, PDB ID: 3OY1)^19^ and glycogen synthase kinase-3*β* (GSK3*β*, PDB ID: 6Y9S)^20^ are particularly compelling due to their synergistic roles in neuronal apoptosis, tau pathology, and neuroinflammation, and have been widely adopted as a benchmark for multi-objective molecular design^15^. For this case study, we initialize EvoSynth with a JNK3-selective inhibitor reported by Shuai et al.^21^, using it as a bioactive scaffold for dual JNK3/GSK3*β* optimization.
- **Ovarian cancer (OC):** One of the most lethal gynecologic malignancies, characterized by late-stage diagnosis, chemoresistance, and poor long-term survival^22^. OC is driven by multiple dysregulated signaling and DNA repair pathways^23^, making it well-suited for multi-target intervention. Phosphoinositide 3-kinase (PI3K, PDB ID: 1E7V)^24^ and poly(ADP-ribose) polymerase 1 (PARP1, PDB ID: 7AAD)^25^ form a complementary therapeutic pair, with preclinical studies showing that their combined inhibition synergistically suppresses OC growth^16^, establishing this dual-target combination as a clinically relevant benchmark. For this case study, we begin with LY-294002^24^, a classical PI3K inhibitor, and optimize it for dual PI3K/PARP1 inhibition.

### 2.1 Efficacy of Latent Evolutionary Optimization with Pharmacological Screening

As detailed in Section 4.2.2, both EvoSynth and MolSculptor implement latent evolutionary optimization in which latent diffusion acts as a stochastic mutation operator and NSGA-II performs multi-objective selection (Fig. 7). To ensure clinical relevance, this optimization is coupled with pharmacological screening, in which candidate molecules are filtered against a comprehensive set of property constraints. In both frameworks, only the binding affinity to therapeutic targets is optimized directly, while all other properties, including SA, QED, and disease-specific ADMET requirements (e.g., permeability of the BBB for AD and toxicity profiles for OC), are enforced as hard feasibility filters. Additional design requirements, such as structural similarity to the lead scaffold and the presence of required substructures, are also treated as constraints. Candidates that fail any filter are discarded regardless of potency. This unified protocol ensured that both methods operated within the same chemical space of molecules with translational potential, allowing observed performance differences to be attributed solely to their evolutionary optimization strategies rather than differences in constraint handling. In fairness, both frameworks were run under identical evolutionary budgets, with fixed population sizes of 128 molecules over 30 generations, matching the setup reported in the MolSculptor study.

We evaluated EvoSynth and MolSculptor from two complementary perspectives. At the efficacy level, we quantified enrichment for potent dual-target binders by analyzing docking scores within the top-quartile subset (Q1), defined as the top 25% of molecules in the population with the highest predicted mean dual-target binding affinity. This focus on Q1 reflects the practical reality of drug discovery, where only a small subset of the most promising candidates advance to experimental validation. At the population quality level, we further assessed the chemical space explored by Q1 through two structural metrics: internal diversity, which measures scaffold variation among top candidates, and similarity to the lead compound, which captures the degree of scaffold exploitation versus exploration.

The results of this comparison are shown in In both case studies (Fig. 1), EvoSynth consistently outperformed MolSculptor in mean dual-target affinity within Q1. In Alzheimer’s disease (JNK3/GSK3*β*), EvoSynth (Contrastive) reached a mean affinity of approximately 11.55 in the final iteration, compared with 11.25 for MolSculptor and 10.61 for EvoSynth (Predictor). In particular, EvoSynth (Contrastive) exhibits a sharp initial improvement during the first few generations, followed by steady incremental gains, reaching MolSculptor’s final affinity level of around 11.28 within 8 generations. A similar trend is observed in OC (PI3K/PARP1), where EvoSynth (Contrastive) achieves a final mean affinity of about 10.68, outperforming both MolSculptor (10.32) and EvoSynth (Predictor, 10.24). Notably, EvoSynth (Contrastive) surpasses MolSculptor’s final affinity as early as generation 12, underscoring its faster convergence. Taken together, these results demonstrate that although the comparison was conducted under a fixed 30-generation budget for consistency with MolSculptor, EvoSynth (Contrastive) achieves superior performance within only a fraction of that budget, highlighting both its effectiveness and efficiency in guiding the evolutionary search.

**Figure 1.**
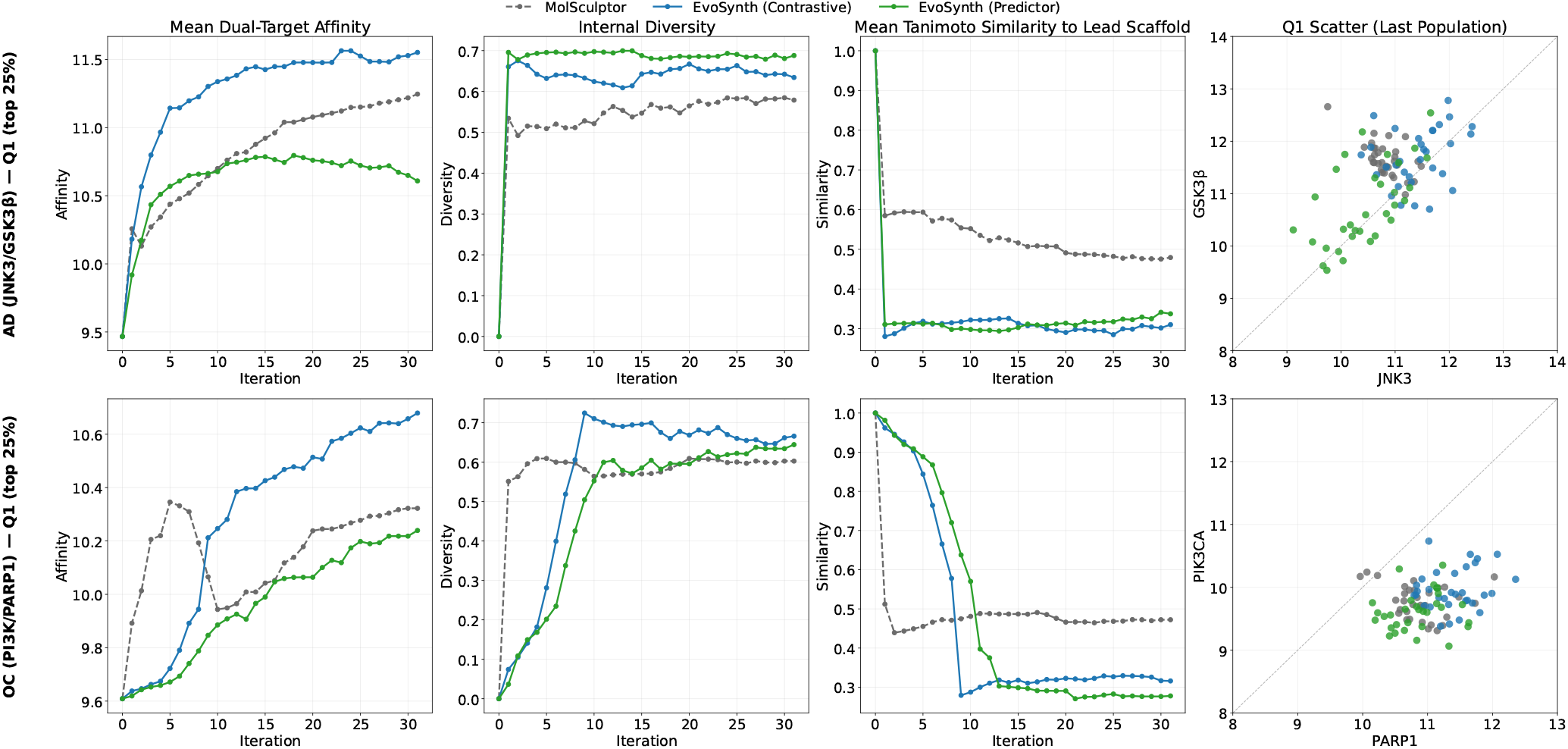
Comparative evaluation of EvoSynth and MolSculptor on dual-target optimization for Alzheimer’s disease (AD; JNK3/GSK3*β*, top row) and ovarian cancer (OC; PARP1/PIK3CA, bottom row). Each row shows optimization trajectories of (i) mean dual-target affinity, (ii) internal diversity, (iii) mean Tanimoto similarity to the lead scaffold, and (iv) Q1 scatter plots of the final populations. All quantitative metrics are computed over the top 25% of molecules (Q1) from each generation to focus on high-performing candidates. Overall, EvoSynth consistently outperforms MolSculptor by achieving higher affinities, greater internal diversity, and substantially lower similarity to the initial lead scaffolds. These results highlight EvoSynth’s ability to generate structurally novel yet pharmacologically potent molecules, effectively balancing exploration and optimization during multi-objective evolution.

In addition to binding affinity, EvoSynth consistently maintains higher scaffold diversity than MolSculptor across both case studies. For Alzheimer’s disease, EvoSynth (Contrastive) achieves a diversity score of around 0.63 compared with about 0.58 for MolSculptor, while also reducing similarity to the seed scaffold from roughly 0.48 (MolSculptor) to 0.31. In OC, a similar trend is observed, where EvoSynth (Contrastive) maintains a diversity of approximately 0.67 versus 0.60 for MolSculptor, accompanied by a drop in lead similarity from about 0.47 to 0.32. These results reflect the richer, score-informed latent space in EvoSynth, which promotes the exploration of chemically novel but pharmacologically relevant regions of chemical space. In contrast, MolSculptor’s reconstruction-oriented latent space yields higher similarity-to-lead values and more homogeneous chemotypes. Although this structural bias maintains stable affinities across its populations, it limits diversity and restricts scaffold innovation. EvoSynth, by contrast, generates structurally distinct yet pharmacologically viable analogues, as reflected in both higher diversity and lower similarity-to-lead values.

The Q1 scatter plots provide a clear view of the joint distribution of the final populations in both targets. In AD (JNK3/GSK3*β*), EvoSynth (Contrastive) produces candidates tightly clustered around (11.20, 11.50) on average, close to the diagonal and indicating consistently high and balanced dual-target binding affinity. MolSculptor, by contrast, centers around (10.80, 11.70), showing stronger GSK3*β* activity but weaker JNK3 binding. EvoSynth (Predictor) remains balanced at (10.40, 10.80) but at a lower absolute affinity than Contrastive. In OC (PI3K/PARP1), EvoSynth (Contrastive) again dominates, with Q1 means of (11.40, 10.00), showing strong PARP1 binding while maintaining moderate PI3K activity. MolSculptor yields (10.90, 9.80), reflecting weaker binding overall, while EvoSynth (Predictor) achieves (10.90, 9.60) at a similarly lower level. Taken together, these scatter plots highlight that EvoSynth (Contrastive) not only enriches for high-affinity binders, but also uncovers more diverse and balanced activity profiles across both targets.

### 2.2 Evaluation of Synthesis-Aware Candidate Prioritization

To assess the practical utility of the molecules generated by EvoSynth and MolSculptor, we applied the synthesis-aware candidate prioritization procedure (Section 4.2.3) to select the top *k* = 10 candidates from their final evolved populations. This procedure formulates candidate selection as a retrosynthetic graph optimization problem (Fig. 8), jointly balancing three objectives: therapeutic reward, initial material cost and reaction reliability. The cutoff of ten candidates reflects a realistic upper bound on the number of molecules that can feasibly be advanced to experimental testing in typical wet-lab pipelines. We evaluated the prioritized sets using four complementary metrics: (i) expected reward, defined as the mean predicted dual-target affinity weighted by the probability of successful synthesis; (ii) number of reaction steps, reflecting overall synthetic effort; (iii) number of unique starting materials, capturing the degree of precursor reuse and potential for batch synthesis; and (iv) total starting material cost, estimating procurement expense. Together, these metrics provide a balanced assessment of both pharmacological promise and synthetic feasibility.

In addition to MolSculptor, we also compare against CMD-GEN^26^, a recently introduced structure-based generative framework that has shown strong performance in dual-target inhibitor design. CMD-GEN operates under a distinct paradigm: instead of evolving molecular populations, it employs a diffusion model to directly generate candidate molecules conditioned on pharmacophore point clouds derived from lead compounds. By aligning pharmacophores across multiple protein pockets, CMD-GEN identifies overlapping regions and samples within these regions to construct dual-target molecules. We include CMD-GEN as a baseline to evaluate our central hypothesis that SPARROW is particularly well suited to EvoSynth’s evolutionary sampling strategy. Because evolutionary sampling naturally generates families of structurally related molecules, it increases the likelihood of shared intermediates and overlapping synthetic routes, which in turn can lead to lower synthesis costs. In contrast, CMD-GEN does not explicitly leverage such structural relationships, making it a useful complementary baseline for assessing the cost-efficiency of synthesis-aware candidate prioritization.

The comparative results are summarized in Fig. 2. In both case studies, EvoSynth (Contrastive) demonstrates the most favorable balance between pharmacological reward and synthetic accessibility. In Alzheimer’s disease (AD; JNK3/GSK3*β*), it achieves the highest expected reward (117.61) while also achieving the lowest synthesis cost (1220 $/g), outperforming both EvoSynth (Predictor, 105.79 reward, 2443 $/g) and MolSculptor (109.49 reward, 3044 $/g). A similar trend is observed in ovarian cancer (OC; PI3K/PARP1), where EvoSynth (Contrastive) reaches 95.74 reward at a moderate cost of 1801 $/g, compared to EvoSynth (Predictor, 92.19 reward, 1277 $/g) and MolSculptor (93.56 reward, 2065 $/g). While the Predictor variant achieves marginally lower synthesis costs in OC, its diminished reward points to reduced potency, underscoring the superior overall balance achieved by the Contrastive approach. By contrast, CMD-GEN yields markedly lower rewards (55.20 in AD and 87.30 in OC) coupled with substantially higher synthesis costs (approximately 3566 $/g in AD and 4590 $/g in OC), highlighting the inherent challenge of generating synthetically accessible molecules in the absence of evolutionary refinement. In terms of synthetic effort, EvoSynth also produces more efficient and convergent synthetic routes. In AD, the Contrastive variant requires 48 steps and 23 starting materials, which is comparable to Predictor (44 steps, 23 materials) but with lower cost and higher reward, while MolSculptor and CMD-GEN generate longer, more fragmented pathways (46 and 56 steps, respectively). Similarly, in OC, EvoSynth variants require only 30–32 steps, substantially fewer than MolSculptor (42) or CMD-GEN (60). The retrosynthetic graphs in Fig. 3 further illustrate these differences: EvoSynth generates sparser, more convergent networks with intermediate reuse and overlapping synthesis routes, whereas CMD-GEN and MolSculptor produce denser, less efficient graphs with specialized precursors and higher synthetic overhead. Collectively, these results highlight the synthesis efficiency and pharmacological advantage of evolutionary design framework.

**Figure 2.**
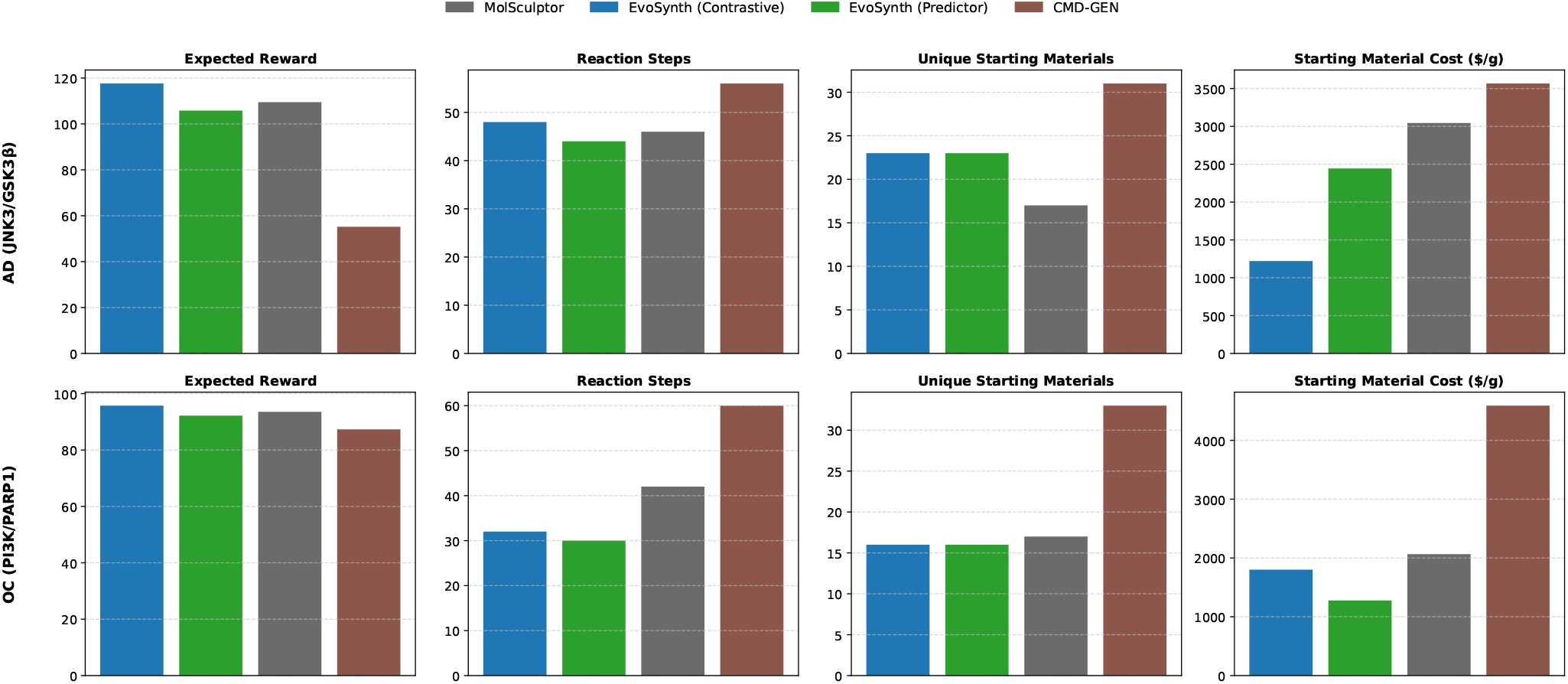
Quantitative evaluation of synthesis-aware candidate prioritization. Bar plots compare EvoSynth (Contrastive), EvoSynth (Predictor), and MolSculptor across two case studies: Alzheimer’s disease (AD, JNK3/GSK3*β*) and ovarian cancer (OC, PI3K/PARP1). Metrics include (i) total expected reward, (ii) number of reaction steps, (iii) number of unique starting materials, and (iv) total starting material cost. Overall, EvoSynth demonstrates the best trade-off across all metrics, achieving the highest expected reward, fewest reaction steps, and lowest synthesis cost, indicating its effectiveness in generating potent yet synthetically accessible candidates.

**Figure 3.**
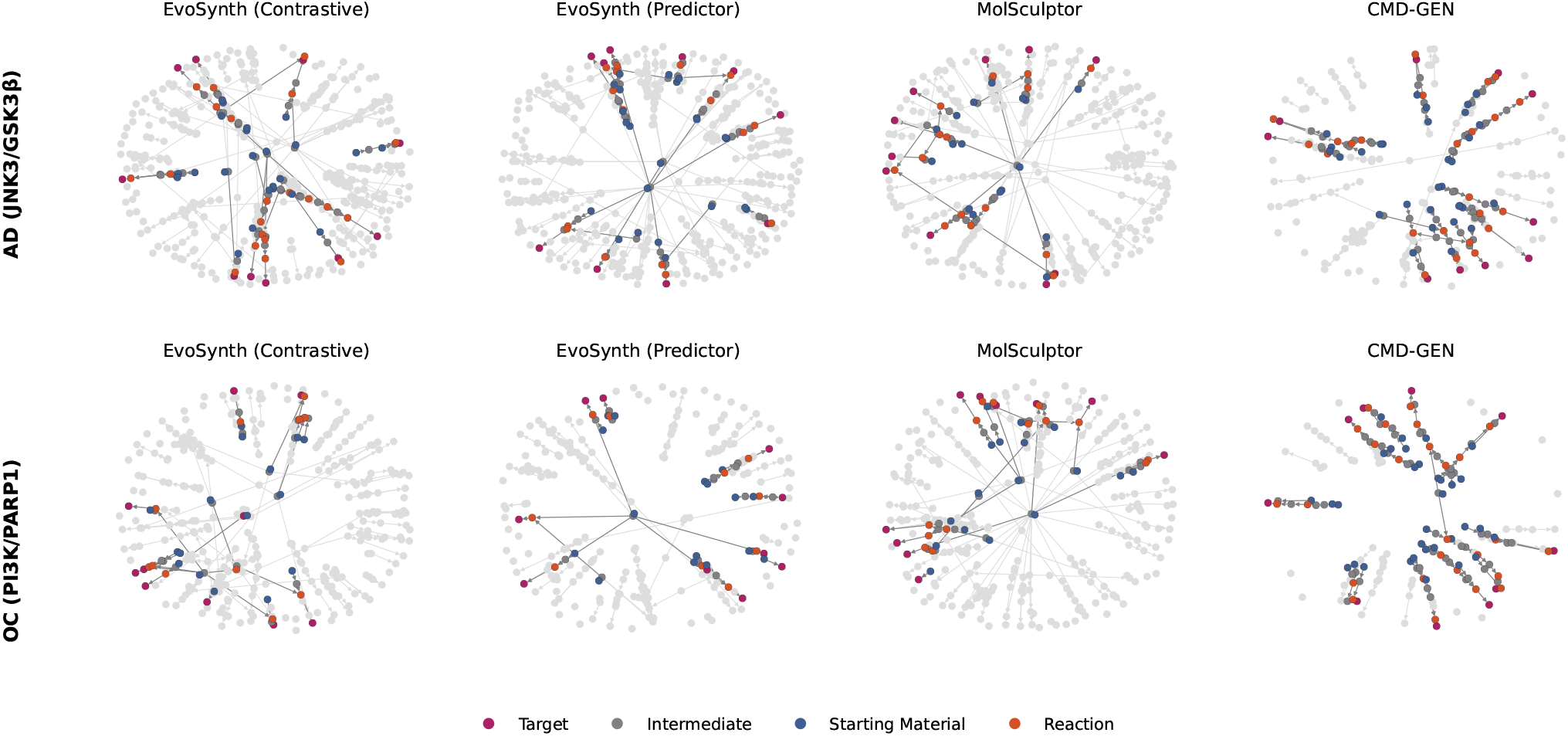
Retrosynthetic networks across methods and case studies. Each panel shows the retrosynthetic graph for the ten selected molecules. Nodes represent therapeutic targets, intermediates, starting materials, and predicted reactions, while highlighted edges denote the synthesis routes chosen as cost-effective and reliable for producing the prioritized candidates. Compared to MolSculptor and CMD-GEN, EvoSynth produces denser, more interconnected synthesis graphs, indicating a higher degree of intermediate sharing and reaction reuse. This structural coherence reflects EvoSynth’s ability to favor synthetically convergent designs-molecules that not only exhibit strong pharmacological potential but are also more amenable to scalable synthesis.

Importantly, EvoSynth achieves this cost reduction without explicitly optimizing for synthetic accessibility. We attribute this effect to two factors: (i) EvoSynth generates a larger pool of candidates with high rewards, giving the SPARROW selector more flexibility to identify molecules that combine potency with cost-effective precursors, and (ii) its greater scaffold diversity promotes overlap across retrosynthetic routes, encouraging the reuse of widely available, inexpensive building blocks. In contrast, MolSculptor’s narrower, lead-focused populations tend to converge on chemotypes linked to costly precursors, increasing the overall synthetic burden. Collectively, these results show that EvoSynth prioritizes candidates that are not only pharmacologically promising, but also more synthetically tractable and better suited for progression to experimental validation, as further illustrated by the representative retrosynthetic routes in Fig. 4.

**Figure 4.**
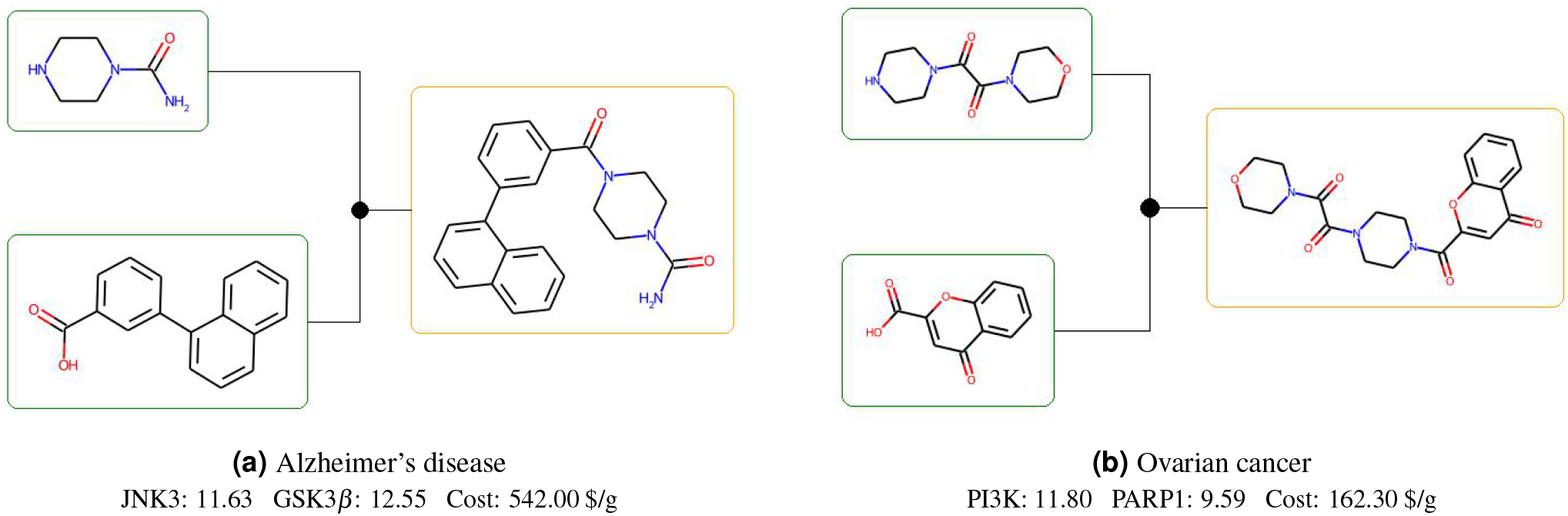
Representative retrosynthetic routes for EvoSynth candidates. Green boxes indicate commercially available starting materials, orange boxes denote synthesized intermediates and the final target molecule, and black dots represent ASKCOS-predicted reaction steps. Binding affinities are reported for each therapeutic target together with the estimated synthetic cost of the starting materials. All candidates generated by EvoSynth are accompanied by such retrosynthetic annotations, ensuring that each proposed molecule is experimentally actionable and can be directly advanced to wet-lab synthesis and validation.

### 2.3 Analysis of Score-Informed Latent Space

A key distinction between EvoSynth and MolSculptor lies in how their latent spaces are constructed. MolSculptor employs a reconstruction-oriented representation trained primarily to capture structural information, ensuring chemical validity but offering limited functional guidance. In contrast, EvoSynth introduces a score-informed latent space that integrates both structural and functional signals, aligning molecular embeddings with predicted multi-target binding profiles so that structural proximity more directly reflects functional similarity, thereby improving the efficiency of evolutionary optimization. EvoSynth’s score-informed latent space was realized through two complementary fine-tuning strategies, contrastive alignment^27^ and predictor-based optimization^28^, as described in Section 4.2.1.

To evaluate the impact of this design, we visualize the latent space landscapes of molecules from the training dataset and compare their organization across models. As shown in Fig. 5, MolSculptor produces a fragmented distribution, with regions of high predicted affinity scattered across the t-SNE projection. This pattern reflects its reconstruction-focused objective, which preserves valid molecular structures but lacks functional alignment. In contrast, EvoSynth exhibits smoother enrichment patterns, where high-affinity molecules form coherent, continuous clusters. This demonstrates that fine-tuning with functional supervision effectively reorganizes the latent space to capture activity gradients, enabling more directed exploration of promising regions.

**Figure 5.**
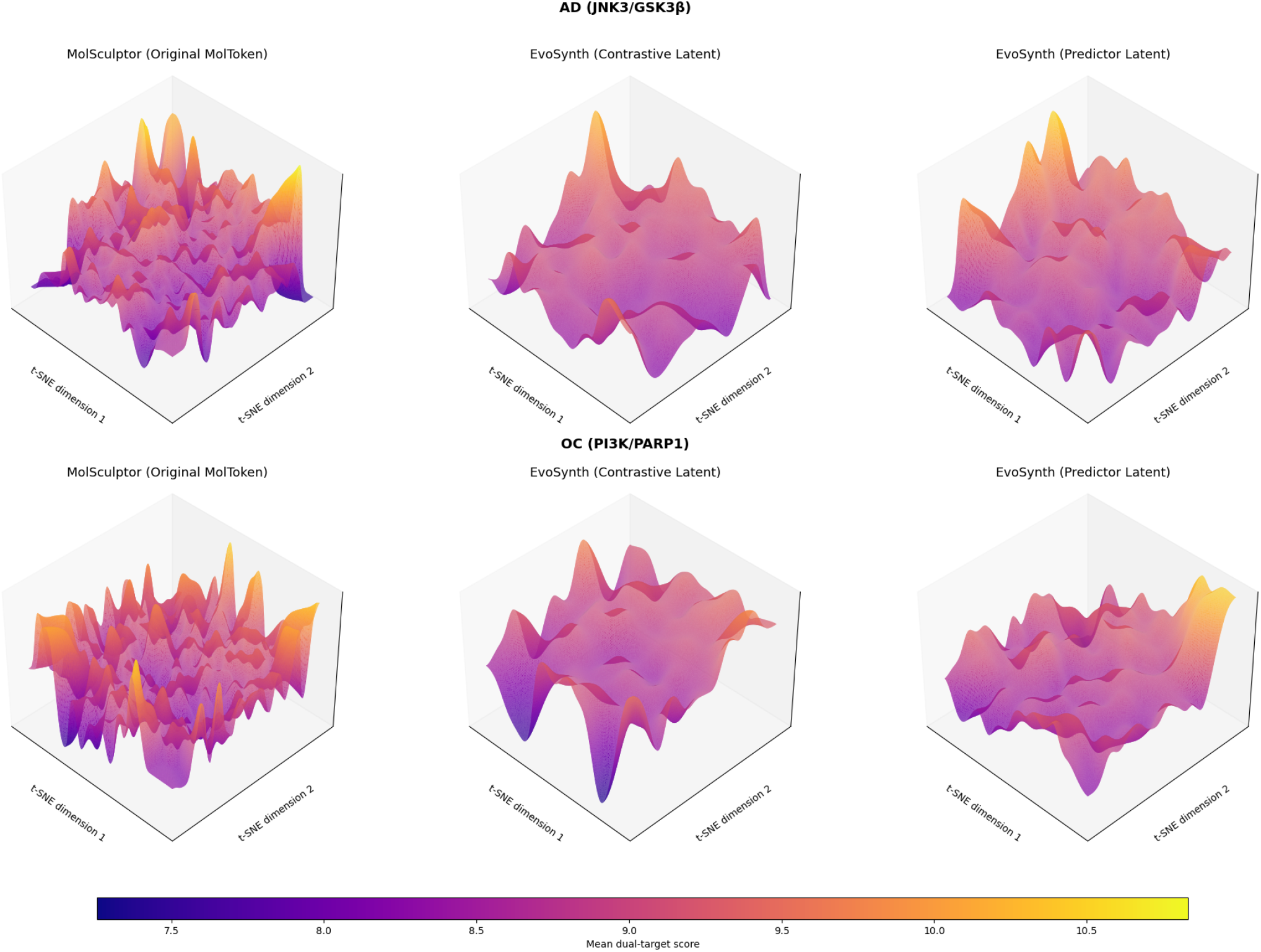
Latent space organization of MolSculptor and EvoSynth across two dual-target case studies. Each surface depicts a 2D t-SNE projection of the latent space colored by mean predicted dual-target binding affinity. Compared to MolSculptor’s fragmented and functionally diffuse landscape, EvoSynth produces smoother, functionally aligned manifolds with coherent high-affinity clusters. This organization enables more directed evolutionary exploration, improving both search efficiency and enrichment of potent scaffolds.

Between the two fine-tuning strategies, the contrastive variant yields a more functionally coherent and stable latent organization. By clustering molecules with similar binding profiles while separating inactive scaffolds, it establishes sharper functional boundaries and clearer enrichment regions. The predictor-based variant, while capable of modeling affinity trends, displays higher variability, likely due to the limited amount of labeled data available in our dual-target setting. Under these conditions, the contrastive objective provides a stronger inductive bias, producing a smoother and more interpretable representation of the chemical space.

### 2.4 Ablation Study on ADMET Property Screening

To further evaluate the role of pharmacological constraints, we conducted an ablation study to assess the impact of ADMET property screening on the evolutionary process of EvoSynth. While the primary optimization objective is binding affinity, we incorporate filters for toxicity and BBB permeability to improve translational relevance. To isolate their contribution, we also executed EvoSynth without enforcing ADMET constraints during optimization. After 30 generations, we retrospectively evaluated the proportion of molecules that met the ADMET criteria, providing information on how these filters shaped the final evolved populations.

For AD, a BBB permeability threshold of 0.7 or above preserved 80.47% of molecules from the unconstrained run still satisfied the requirement, whereas for OC, a ClinTox cutoff of 0.3 or below yielded yielded 67.19% compliance. Surprisingly, removing the ADMET constraints did *not* accelerate progress. As shown in Fig. 6, the unconstrained runs converged more slowly and achieved a lower final affinity, measured as the mean dual-target affinity within Q1 in the final iteration (11.33 vs.

**Figure 6.**
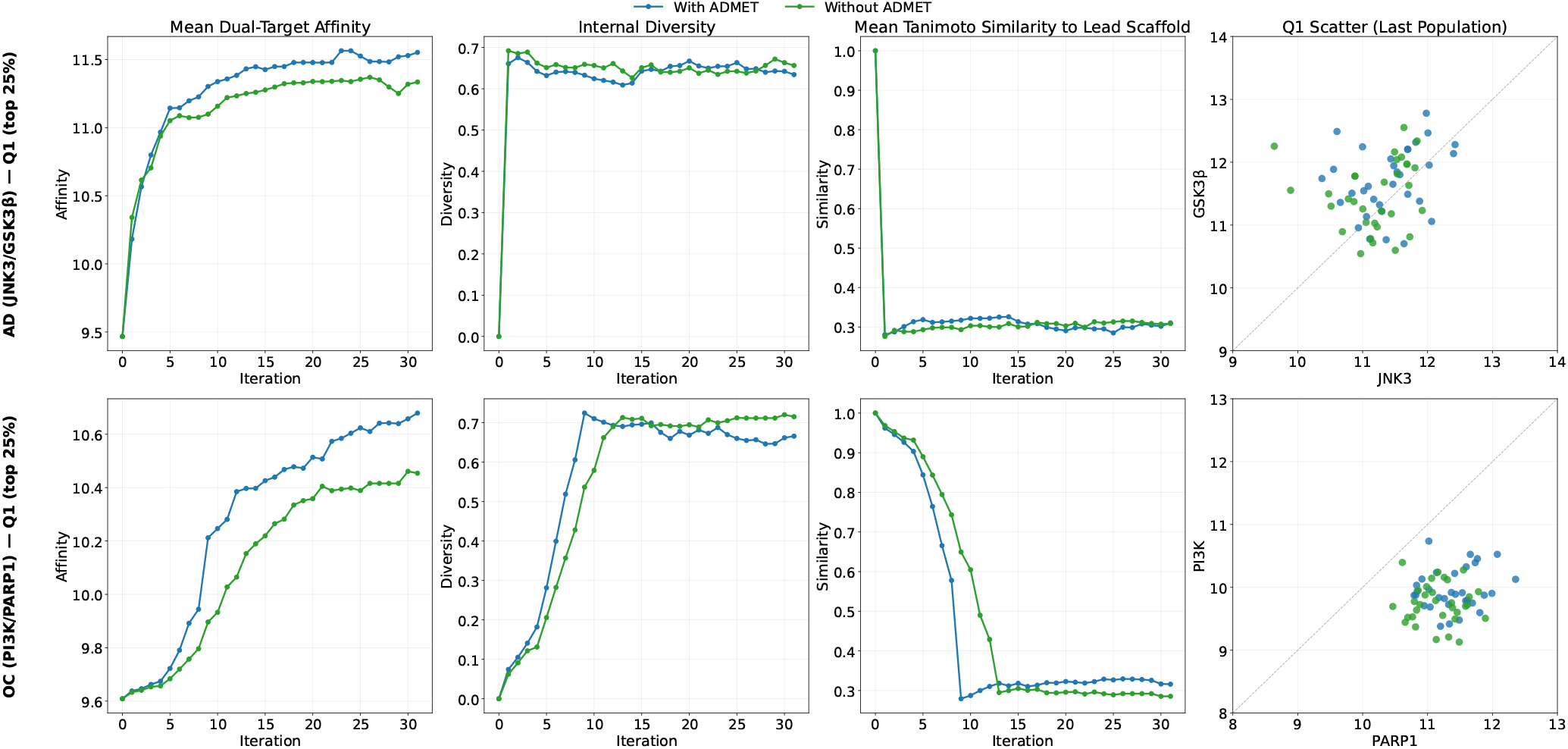
Ablation study on ADMET property screening using EvoSynth (Contrastive). Comparison of optimization trajectories with (blue) and without (green) ADMET constraints for AD (top) and OC (bottom). Incorporating ADMET screening consistently improved convergence and produced more pharmacologically viable molecules, demonstrating that these constraints serve as an effective inductive bias rather than a limiting factor in EvoSynth’s optimization process.

11.55 in AD; 10.45 vs. 10.68 in OC), compared to the ADMET-constrained runs. These findings suggest that ADMET screening provides a beneficial inductive bias: by pruning pharmacologically unfavorable candidates during the search, EvoSynth is steered toward regions of chemical space that are both clinically viable and more amenable to optimization. Thus, rather than restricting exploration, ADMET constraints enhance both the efficiency and the effectiveness of the evolutionary search process.

## 3 Discussion

The central challenge addressed in this work is the design of molecules with balanced activity across multiple protein targets, a goal where most generative frameworks fall short. EvoSynth advances this objective through three key contributions. **(1) Functional alignment through score-informed latent representation:** By embedding affinity signals into the molecular latent space, EvoSynth tightly couples structural and functional features, yielding more organized representations, diverse scaffolds, and stronger enrichment for high-affinity candidates. **(2) Integrated optimization under pharmacological and synthetic constraints:** Beyond affinity optimization, EvoSynth incorporates pharmacological filters and retrosynthetic feasibility during design, improving convergence and producing candidates that are both pharmacologically viable and synthetically accessible. **(3) Synthesis-aware prioritization via SPARROW integration:** To ensure practical feasibility, EvoSynth employs SPARROW to jointly optimize therapeutic reward, reaction cost, and route reliability. This integration identifies high-reward, low-cost molecules and promotes structural reuse and route convergence, bridging functional optimization with real-world synthetic constraints. Together, these contributions position EvoSynth as a unified framework for multi-objective molecular design that balances potency, accessibility, and practicality.

Despite these advances, several limitations suggest promising directions for future research. **(1) Variability from multimodal latent alignment:** Integrating structural and functional signals enriches exploration but introduces variability in predicted affinities, as reconstruction objectives are smooth whereas functional supervision is noisier. This could be mitigated by adaptive weighting between the structural and functional losses or by incorporating uncertainty-aware training to stabilize functional gradients. **(2) Dependence on docking-derived supervision:** The current reliance on docking-based affinity labels ties performance to docking accuracy and target coverage. Future work could replace or augment these labels with experimental or high-confidence binding data, or employ transfer learning from large-scale protein–ligand interaction datasets to improve generalization. **(3) Post-hoc treatment of synthetic feasibility:** SPARROW is presently applied only after evolution, allowing some infeasible yet high-reward molecules to persist. Incorporating synthesis-aware objectives directly into the optimization loop, using machine learning surrogates trained on ASKCOS-labeled data, would enable real-time trade-offs between potency and practicality. **(4) Limited scope to dual-target optimization:** This study focuses on dual-target design as a controlled proof of concept. Extending EvoSynth to polypharmacological optimization, including simultaneous multi-target efficacy and off-target penalty modeling, would enhance clinical relevance and support the design of more effective and selective therapeutics.

## 4 Methods

### 4.1 Problem Formulation

EvoSynth aims to generate multi-target therapeutics that simultaneously maximize predicted binding affinity and satisfy pharmacological and synthetic constraints. We define a therapeutic context as the pair (*{T*_1_, …, *T*_*m*_*},𝒦*), where 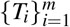 is the set of molecular targets and 𝒦 denotes disease-specific requirements (for example, for neurological indications 𝒦 may include thresholds of permeability of the BBB). We represent a candidate molecule by *x* ∈ *𝒳* (the chemical space) and by **z** ∈ *𝒵* its continuous latent embedding under a pre-trained encoder *E* : *𝒳* → *𝒵* with decoder *D* : *𝒵* → *𝒳* . The design objective is a constrained multi-objective optimization problem over latent space:

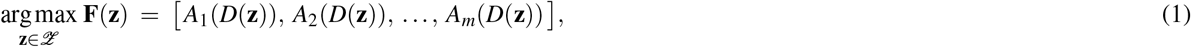

subject to

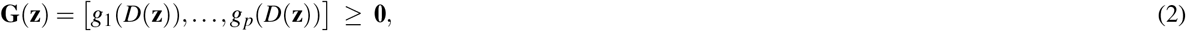

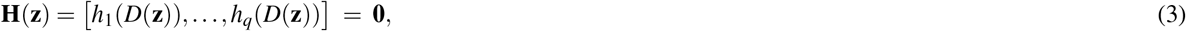

where *A*_*i*_(*x*) denotes the predicted affinity (or reward) for target *T*_*i*_; **G** defines inequality constraints that enforce predefined thresholds on key pharmacological properties (e.g., SA, QED, ADMET-related metrics, and scaffold similarity to the lead compound); and **H** specifies equality constraints that ensure the exact preservation of designated substructures.

To incorporate synthesis-aware trade-offs, we augment the objective vector with synthesis cost and reliability terms derived from retrosynthetic analysis:

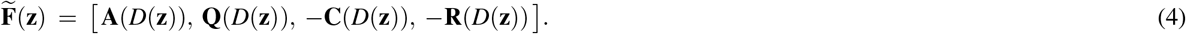

where **Q** denotes pharmacological desirability scores (higher is better), **C** the estimated retrosynthetic material cost, and **R** the synthesis risk (lower is better). Negative signs in Equation (4) place cost and risk in a maximization formulation.

EvoSynth addresses the optimization problem in Equations (1)–(3) through three core components that align with the following subsections: (i) *score-informed latent space learning*, which organizes the latent space *𝒵* so that local geometry captures multi-target functional similarity while enabling stochastic mutation via latent diffusion; (ii) *evolutionary optimization of multi-target affinity*, which applies NSGA-II on the augmented objective 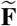 to approximate Pareto-optimal candidates under pharmacological constraints **G** and structural constraints **H**; and (iii) *synthesis-aware candidate prioritization*, which leverages retrosynthetic graph optimization (SPARROW) to jointly balance therapeutic reward, material cost, and reaction reliability. Together, these stages yield candidates that are both pharmacologically promising and synthetically feasible.

### 4.2 Core Framework Components

#### 4.2.1 Score-Informed Latent Space Learning

Given a two-dimensional molecular graph *x*, the MolToken encoder produces an initial latent representation **z** = E(*x*). We define its normalized form as 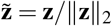, which is projected onto the unit hypersphere and used in contrastive and uniformity objectives. The fine-tuning objective integrates structural reconstruction, latent regularization, and functional alignment:

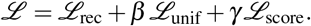

Here, *ℒ*_rec_ enforces faithful molecular reconstruction, *ℒ*_unif_ regularizes the latent distribution for uniformity, and *ℒ*_score_ introduces task-specific functional alignment.

##### Reconstruction Loss

The reconstruction term *ℒ*_rec_ is a cross-entropy loss between predicted and ground-truth molecular tokens, ensuring that the encoder–decoder pathway accurately recovers the input structure:

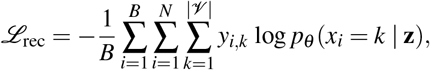

where *B* denotes the batch size, *N* is the length of the token sequence, *𝒱* the vocabulary, *y*_*i,k*_ a one-hot indicator for the token *x*_*i*_, and *p*_*θ*_ (*x*_*i*_ = *k* | **z**) the likelihood of the decoder.

##### Uniformity Loss

To prevent representational collapse, the uniformity objective *ℒ*_unif_ encourages normalized latents 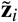 to be evenly distributed on the unit hypersphere, following SimCLR^29^:

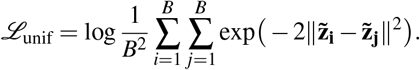

This regularizer promotes a uniform distribution of latent vectors, thereby enhancing the exploration of the latent space.

##### Score-aware Loss

Functional alignment is imposed through a score-aware term *ℒ*_score_, instantiated in two complementary forms:

i. *Contrastive alignment*. An InfoNCE-style objective aligns latent representations according to binding profile similarity, with the positive index *p*(*i*) for each molecule *i* defined as the index of the batch member with the closest predicted binding affinity value (excluding *i* itself):

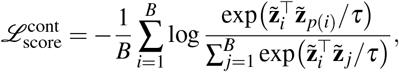

where *τ* is a temperature hyperparameter.
ii. *Predictor-based regression*. Alternatively, a regression head *f*_*θ*_ maps latent vectors to binding scores by mean squared error:

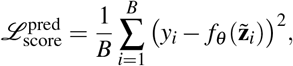

where *y*_*i*_ is the experimental binding score. Together, these objectives balance structural fidelity, representational robustness, and functional alignment within the latent space. The resulting score-informed latent vectors **z** are then fixed and used as training data for the subsequent diffusion model.

#### 4.2.2 Evolutionary Optimization of Multi-Target Affinity

##### In Silico Mutagenesis via Latent Diffusion

After training, the encoder and decoder of the MolToken autoencoder are frozen, producing a stable latent space *𝒵* that provides the substrate for molecular evolution. In this space, mutational exploration is implemented through a diffusion process that serves as a stochastic mutation operator (Fig. 7A).

**Figure 7.**
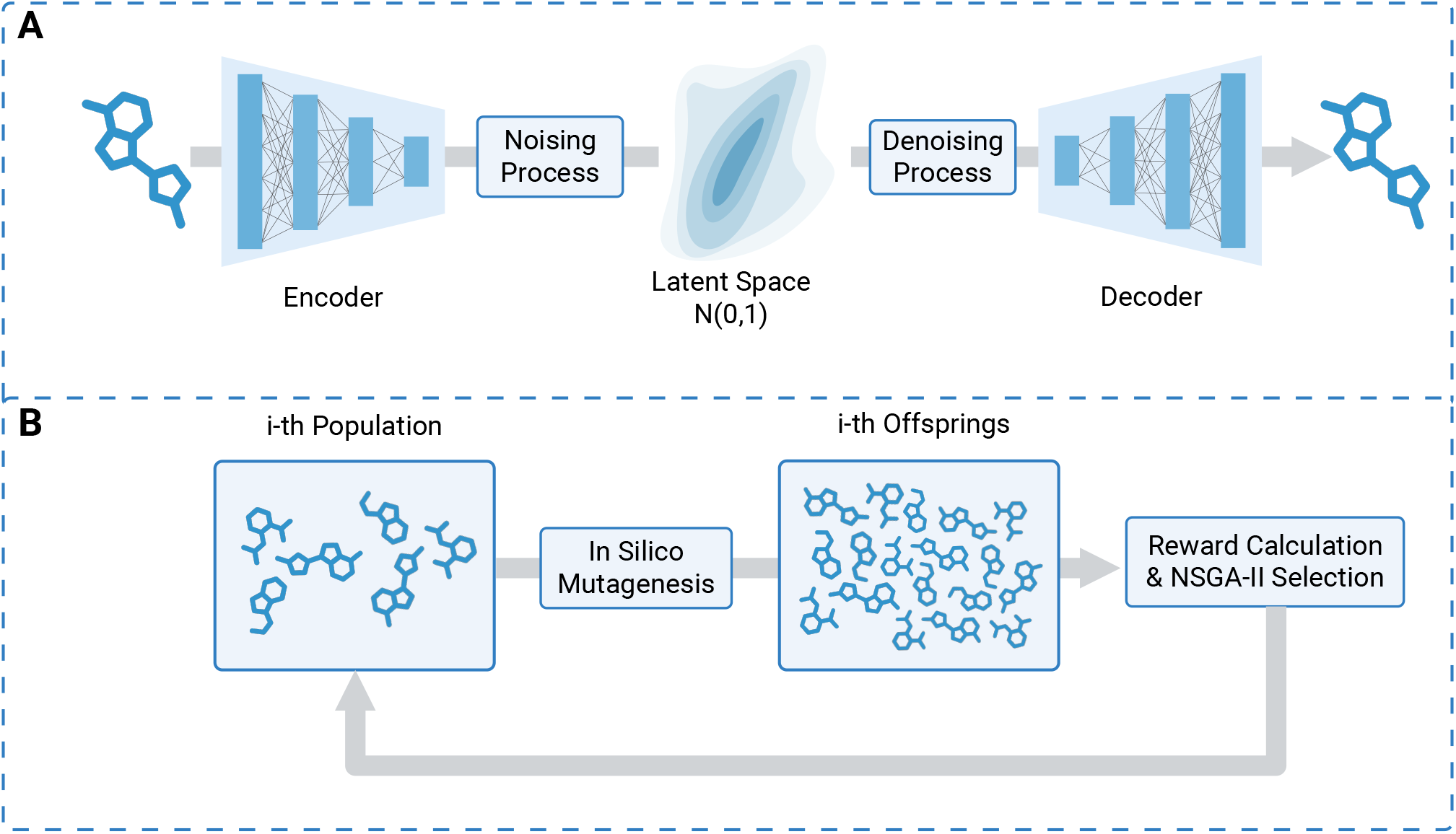
Overview of the latent evolution framework for multi-target affinity optimization. (**A**) In silico mutagenesis in the score-informed latent space. The fine-tuned MolToken autoencoder encodes molecules into a latent representation capturing both structural and functional features. Latent diffusion, acting as a mutation operator, perturbs points sampled from the standard normal prior 𝒩(0, 1) through a noise–denoise process to generate structural variants, which are then decoded into chemically valid candidates. (**B**) Evolutionary optimization loop. An initial population is iteratively evolved by generating offspring through the noise–denoise cycle, followed by reward evaluation based on predicted multi-target affinity and pharmacological filters. The top-scoring molecules are selected to seed the next generation, driving exploration toward promising regions of chemical space.

**Figure 8.**
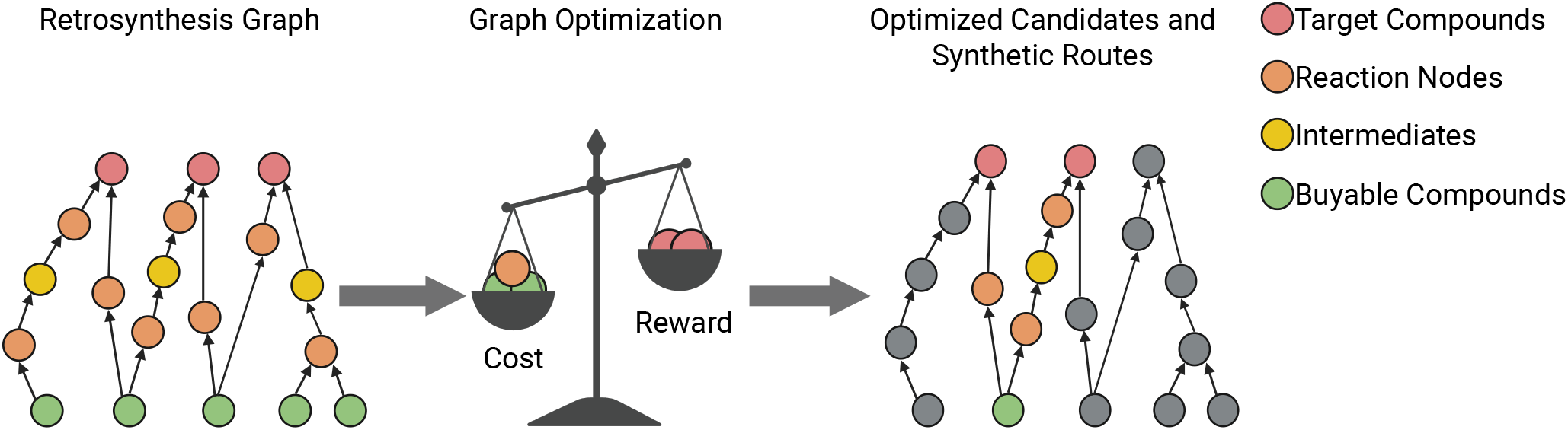
Synthesis-aware candidate prioritization via retrosynthetic graph optimization. The input consists of a retrosynthetic graph that encodes candidate compounds, reactions, and purchasable precursors. The optimization process jointly balances therapeutic reward (predicted affinities), synthetic cost, and reaction reliability. The output is a prioritized subset of candidate compounds accompanied by feasible synthetic routes.

##### Training Objective

The diffusion model is trained to predict Gaussian noise added to the latent vectors, with the *ε*-prediction loss guiding the denoiser to accurately estimate these perturbations:

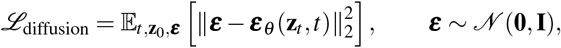

where the timestep *t* is provided to the DiT backbone via adaptive layer normalization (adaLN), enabling time-dependent denoising behavior.

Having established this training objective, we next describe how the diffusion process is applied at inference time to perform in silico mutagenesis.

##### Initialization

Given a lead compound *x*_0_ ∈ *𝒳* represented as a molecular graph, the MolToken encoder maps its structure to a score-informed latent vector **z**_0_. This vector is normalized by its ℓ_2_-norm to yield 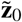, which serves as the starting point for diffusion.

##### Forward Mutation

To generate structurally diverse yet potentially bioactive variants, the latent vector 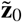 is progressively perturbed through the forward diffusion process:

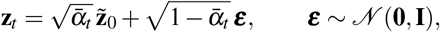

where 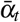 follows a predefined noise schedule and *t* ∈ *{*1,…, *T}*.

##### Reverse Refinement

The trained denoiser then iteratively reconstructs **z**_*t*−1_ from **z**_*t*_:

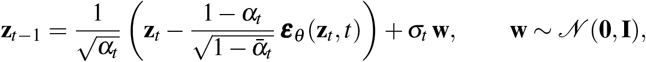

where *α*_*t*_ and *σ*_*t*_ are schedule parameters. Repeated over *T* steps, this process progressively refines the noisy trajectory back to a valid latent representation.

##### Reconstruction

After *T* denoising steps, the model yields a mutated latent vector 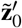, corresponding to a structural analogue of the original compound. The pre-trained decoder subsequently maps this latent back to chemical space, producing a SMILES sequence 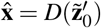 that preserves chemical validity.

Together, this training–inference loop enables in silico mutagenesis directly in the ligand latent space, capturing structural variations and chemical optimization while maintaining plausibility.

#### Computation of Reward and Pharmacological Properties

The reward signal for optimization is defined by the docking-derived binding affinity to the therapeutic targets of interest, estimated using the DSDP framework^30^. Since docking scores are reported on a minimization scale (lower is better), we take their negative values so that higher rewards correspond to stronger binding. Formally, for each latent vector **z** ∈ *𝒵*, the decoded molecule is *D*(**z**), and the reward for the target *T*_*i*_ is

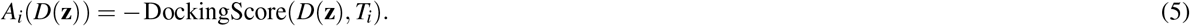

In addition to affinity, candidate molecules are screened for pharmacological and design constraints. Synthetic accessibility (SA), drug-likeness (QED), and scaffold similarity to the lead compound are computed using RDKit^31^, with similarity quantified as the Tanimoto coefficient between Morgan fingerprints (radius = 2). Required substructures are verified through SMARTS-based substructure matching to ensure exact preservation of designated fragments. These properties define constraint functions consistent with Equations (2)–(3):

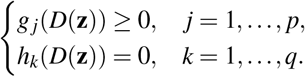

##### Multi-Objective Selection under Constraints

Each property *p* is compared with a predefined threshold *τ*_*p*_, resulting in a binary feasibility indicator:

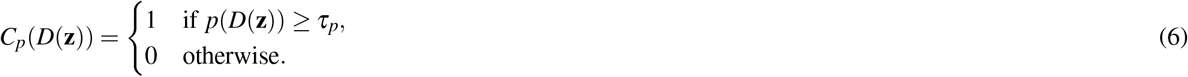

A candidate is feasible only if all constraints are satisfied simultaneously *C*(*D*(**z**)) = ∏_*p*_ *C*_*p*_(*D*(**z**)) = 1. The feasible region in latent space is therefore *ℒ*_feasible_ = **z** *ℒ C*(*D*(**z**)) = 1 within this feasible set, and EvoSynth optimizes the multi-target affinity vector:

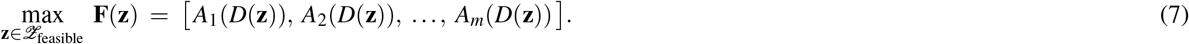

We utilize NSGA-II^32^ to approximate the Pareto front of **F**(**z**), generating a diverse set of Pareto-optimal molecules that balance affinities between targets. These solutions are reintroduced iteratively as the parent population for the next generation, ensuring progressive refinement of potency and diversity while preserving pharmacological and synthetic feasibility.

#### 4.2.3 Synthesis-Aware Candidate Prioritization

Drawing on the SPARROW framework^14^, we formulate compound selection as a multi-objective graph optimization problem on retrosynthetic networks, as illustrated in Fig. 8. The retrosynthetic graph is a directed bipartite structure comprising compound and reaction nodes, with edges representing reactant–product relationships. Commercially available precursors are incorporated as terminal nodes, connected via zero-cost dummy reactions, while reaction nodes are assigned weights based on predicted success probabilities. These graphs are automatically generated using ASKCOS^12^, which provides condition recommendations and forward-predicted reaction likelihoods. The optimization involves two sets of binary decision variables: compound and reaction selection, subject to constraints that ensure chemically valid acyclic synthesis routes originating from purchasable precursors, allowing for flexible choices between purchasing compounds or synthesizing them from cost-effective starting materials.

We employ a scalarized linear objective that balances three key criteria: (i) maximize the cumulative therapeutic reward of selected compounds, quantified as predicted affinities **A** to target proteins; (ii) minimize the total cost of synthesis **C** of required starting materials, sourced from the ChemSpace API^33^ based on availability and pricing; and (iii) reduce penalties for reactions with low reliability **R**, weighted by predicted success probabilities. Adjustable weighting factors enable systematic trade-off analysis among potency, cost, and synthetic reliability, ensuring that prioritized candidates are both pharmacologically promising and synthetically feasible. The optimization problem is expressed as:

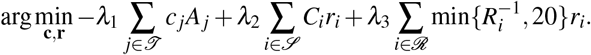

Here **c** and **r** are binary selection vectors for compounds and reactions; *𝒯, 𝒮, ℛ* index the sets of candidate compounds, available starting materials and feasible reactions, respectively. For compound *j, A*_*j*_ denotes the predicted affinity (therapeutic reward); for precursor *i, C*_*i*_ is the estimated cost (sourced from the ChemSpace API^33^); and *R*_*i*_ is the predicted reliability of reaction *i*, with min 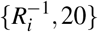 capping extreme penalties. The non-negative weights *λ*_1_, *λ*_2_, *λ*_3_ are used to normalise and tune trade-offs between potency, cost and synthetic reliability.

### 4.3 Materials and Evaluation

#### Materials

The scarcity of case-specific docking data, as noted in the MolSculptor framework, limits direct supervised fine-tuning. To address this, we constructed case-specific datasets for AD and OC case studies. For each, we selected the top 100,000 molecules most similar to the lead compound, based on the Tanimoto similarity of Morgan’s fingerprints, from the MolToken pretraining corpus. This similarity-enriched subset focused on docking computations on chemically relevant candidates, ensuring computational tractability. The molecules were docked against relevant protein targets using DSDP^30^, with the binding affinity scores serving as functional labels for fine-tuning. These docking datasets will be publicly released to support future multi-target drug discovery research.

#### Evaluation Metrics

Binding affinity, reported as predicted docking-derived scores to dual targets, serves as the primary performance metric in our analysis. In addition, we report three supporting metrics that require explicit computation:

i. *Internal diversity*, computed as the mean pairwise Tanimoto distance between Morgan fingerprints within the final population *𝒫*:

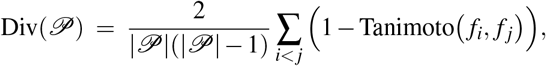

where *f*_*i*_ and *f* _*j*_ denote the Morgan fingerprints of molecules *i* and *j*.
ii. *Mean similarity to lead scaffold*, calculated as the average Tanimoto similarity between each generated molecule and the case-specific seed scaffold *f*_lead_:

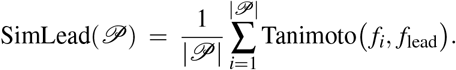
iii. *Expected reward*, defined as the mean predicted dual-target affinity weighted by the probability of successful synthesis:

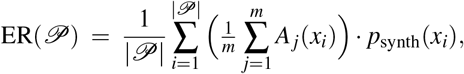

where *A*_*j*_(*x*_*i*_) is the predicted affinity of molecule *x*_*i*_ to target *T*_*j*_, *m* is the number of targets, and *p*_synth_(*x*_*i*_) is the estimated synthesis probability from retrosynthetic analysis.

We also report synthesis-aware metrics, including the total number of reaction steps, the number of unique starting materials, and the cumulative cost of starting materials, all derived directly from retrosynthetic graph analysis.

## Data Availability

All datasets used in this study are derived from publicly accessible sources. Processed data and docking results necessary to reproduce the findings are publicly available at https://github.com/HySonLab/EvoSynth.

## Code Availability

The source code for EvoSynth, including inference pipelines, and retrosynthesis analysis tools, is publicly available at https://github.com/HySonLab/EvoSynth. Pretrained EvoSynth checkpoints required for inference are publicly available on Zenodo at https://zenodo.org/record/17351094.

## Author Contributions Statement

T.S.H. and V.T.D.N. conceived the research idea. V.T.D.N. developed the EvoSynth framework, implemented the experiments, and performed data analysis. P.P. contributed to conceptual refinement, evaluation design and code optimization. V.T.D.N. and

P.P. jointly wrote the initial manuscript draft. T.S.H. supervised the project and provided critical feedback and revisions. All authors discussed the results and approved the final manuscript.

## References

1. Ramsay, R. R., Popovic-Nikolic, M. R., Nikolic, K., Uliassi, E. & Bolognesi, M. L. A perspective on multi-target drug discovery and design for complex diseases. Clin. translational medicine 7, 3 (2018).

2. Halli-Tierney, A. D., Scarbrough, C. & Carroll, D. Polypharmacy: evaluating risks and deprescribing. Am. family physician 100, 32–38 (2019).

3. Makhoba, X. H., Viegas Jr, C., Mosa, R. A., Viegas, F. P. & Pooe, O. J. Potential impact of the multi-target drug approach in the treatment of some complex diseases. Drug design, development therapy 3235–3249 (2020).

4. Katsoulaki, E.-E., Dimopoulos, D. & Hadjipavlou-Litina, D. Multitarget compounds designed for alzheimer, parkinson, and huntington neurodegeneration diseases. Pharmaceuticals 18, 831 (2025).

5. Doostmohammadi, A., Jooya, H., Ghorbanian, K., Gohari, S. & Dadashpour, M. Potentials and future perspectives of multi-target drugs in cancer treatment: the next generation anti-cancer agents. Cell Commun. Signal. 22, 228 (2024).

6. Tang, X. et al. A survey of generative ai for de novo drug design: new frontiers in molecule and protein generation. Briefings Bioinforma. 25 (2024).

7. Li, Y., Lin, X., Hao, Y., Zhang, J. & Gao, Y. Q. Molsculptor: a diffusion-evolution framework for multi-site inhibitor design. ChemRxiv DOI: 10.26434/chemrxiv-2025-v4758 (2025).

8. Ertl, P. & Schuffenhauer, A. Estimation of synthetic accessibility score of drug-like molecules based on molecular complexity and fragment contributions. J. cheminformatics 1, 8 (2009).

9. Bickerton, G. R., Paolini, G. V., Besnard, J., Muresan, S. & Hopkins, A. L. Quantifying the chemical beauty of drugs. Nat. chemistry 4, 90–98 (2012).

10. Swanson, K. et al. Generative ai for designing and validating easily synthesizable and structurally novel antibiotics. Nat. machine intelligence 6, 338–353 (2024).

11. Koziarski, M. et al. Rgfn: Synthesizable molecular generation using gflownets. Adv. Neural Inf. Process. Syst. 37, 46908–46955 (2024).

12. Coley, C. W., Barzilay, R., Jaakkola, T. S., Green, W. H. & Jensen, K. F. Prediction of organic reaction outcomes using machine learning. ACS central science 3, 434–443 (2017).

13. Genheden, S. et al. Aizynthfinder: a fast, robust and flexible open-source software for retrosynthetic planning. J. cheminformatics 12, 70 (2020).

14. Fromer, J. C. & Coley, C. W. An algorithmic framework for synthetic cost-aware decision making in molecular design. Nat. Comput. Sci. 4, 440–450 (2024).

15. Kim, H. et al. Targeting jnk3 for alzheimer’s disease: Design and synthesis of novel inhibitors with aryl group diversity utilizing wide pocket. Eur. J. Medicinal Chem. 285, 117209 (2025).

16. Wang, D. et al. Combined inhibition of pi3k and parp is effective in the treatment of ovarian cancer cells with wild-type pik3ca genes. Gynecol. oncology 142, 548–556 (2016).

17. Scheltens, P. et al. Alzheimer’s disease. The Lancet 397, 1577–1590 (2021).

18. Gong, C.-X., Liu, F. & Iqbal, K. Multifactorial hypothesis and multi-targets for alzheimer’s disease. J. Alzheimer’s Dis. 64, S107–S117 (2018).

19. Probst, G. D. et al. Highly selective c-jun n-terminal kinase (jnk) 2 and 3 inhibitors with in vitro cns-like pharmacokinetic properties prevent neurodegeneration. Bioorganic & medicinal chemistry letters 21, 315–319 (2011).

20. Buonfiglio, R. et al. Discovery of novel imidazopyridine gsk-3β inhibitors supported by computational approaches. Molecules 25, 2163 (2020).

21. Shuai, W. et al. Discovery of novel indazole chemotypes as isoform-selective jnk3 inhibitors for the treatment of parkinson’s disease. J. Medicinal Chem. 66, 1273–1300 (2023).

22. Jayson, G. C., Kohn, E. C., Kitchener, H. C. & Ledermann, J. A. Ovarian cancer. The lancet 384, 1376–1388 (2014).

23. Murawski, M., Jagodziński, A., Bielawska-Pohl, A. & Klimczak, A. Complexity of the genetic background of oncogenesis in ovarian cancer—genetic instability and clinical implications. Cells 13, 345 (2024).

24. Walker, E. H. et al. Structural determinants of phosphoinositide 3-kinase inhibition by wortmannin, ly294002, quercetin, myricetin, and staurosporine. Mol. cell 6, 909–919 (2000).

25. Ogden, T. E. et al. Dynamics of the hd regulatory subdomain of parp-1; substrate access and allostery in parp activation and inhibition. Nucleic acids research 49, 2266–2288 (2021).

26. Zou, Y. et al. A structure-based framework for selective inhibitor design and optimization. Commun. Biol. 8, 422 (2025).

27. Oord, A. v. d., Li, Y. & Vinyals, O. Representation learning with contrastive predictive coding. arXiv preprint arXiv:1807.03748 (2018).

28. Castro, E. et al. Transformer-based protein generation with regularized latent space optimization. Nat. Mach. Intell. 4, 840–851 (2022).

29. Chen, T., Kornblith, S., Norouzi, M. & Hinton, G. A simple framework for contrastive learning of visual representations. In International conference on machine learning, 1597–1607 (PmLR, 2020).

30. Huang, Y. et al. Dsdp: A blind docking strategy accelerated by gpus. J. chemical information modeling 63, 4355–4363 (2023).

31. RDKit — rdkit.org. https://www.rdkit.org/. [Accessed 29-08-2025].

32. Deb, K., Pratap, A., Agarwal, S. & Meyarivan, T. A fast and elitist multiobjective genetic algorithm: Nsga-ii. IEEE transactions on evolutionary computation 6, 182–197 (2002).

33. Chemspace Services: Compound Sourcing and Procurement, Hit Discovery, Molecular Docking, Custom Synt — chemspace.com. https://chem-space.com/services. [Accessed 12-08-2025].

